# Phylogenetic placement of short reads without sequence alignment

**DOI:** 10.1101/2020.10.19.344986

**Authors:** Matthias Blanke, Burkhard Morgenstern

## Abstract

Phylogenetic placement is the task of placing a query sequence of unknown taxonomic origin into a given phylogenetic tree of a set of reference sequences. Several approaches to phylogenetic placement have been proposed in recent years. The most accurate of them need a multiple alignment of the reference sequences as input. Most of them also need alignments of the query sequences to the multiple alignment of the reference sequences. A major field of application of phylogenetic placement is taxonomic read assignment in metagenomics.

Herein, we propose *App-SpaM*, an efficient alignment-free algorithm for phylogenetic placement of short sequencing reads on a tree of a set of reference genomes. *App-SpaM* is based on the *Filtered Spaced Word Matches* approach that we previously developed. Unlike other methods, our approach neither requires a multiple alignment of the reference genomes, nor alignments of the queries to the reference sequences. Moreover, *App-SpaM* works not only on assembled reference genomes, but can also take reference taxa as input for which only unassembled read sequences are available.

The quality of the results achieved with *App-SpaM* is comparable to the best available approaches to phylogenetic placement. However, since *App-SpaM* is not based on sequence alignment, it is between one and two orders of magnitude faster than those existing methods.

## Background

Phylogeny reconstruction is a fundamental field of research in bioinformatics [12]. Here, the basic task is to reconstruct a phylogenetic tree for a set of nucleic-acid or protein sequences, representing their evolutionary history. With the amount of sequence data now available, and with the existing knowledge about phylogeny, however, it is often neither necessary nor desirable to construct phylogenetic trees from scratch. If a reliable phylogenetic tree is already known for a subset of the input sequences, then it is more reliable and more efficient to find the position of the remaining sequences within this existing tree. This procedure is called *phylogenetic placement*; a number of algorithms have been proposed for this task during the last years [25, 28, 7, 3].

In metagenomics studies, DNA from environmental samples is directly sequenced to study microbial communities, without cultivating the sequenced microbes [9, 35]. A fundamental step in metagenomics data analysis is to assign sequencing reads from an environmental sample to known taxa. This task is known as *taxonomic read assignment*. Numerous approaches have been proposed for taxonomic read assignment, *e.g*. [34, 27, 33, 10] see [6] for review.

Earlier approaches to taxonomic read assignment relied on database search programs such as *BLAST* [1]. This idea has been implemented, for example in the widely-used program *MEGAN* [15, 17, 16]. Since using *BLAST* became too slow for the large sets of reads produced in metagenomic projects, k-mer based programs have been developed, such as *TETRA* [37, 36], *Kraken* [38, 39] or *CLARK-S* [31]. This has been generalized to using so-called *spaced-words* instead of contiguous k-mers [8, 14]. Most of the above methods classify reads at predefined taxonomic levels, *e.g*. at the *species* or *genus* level. This way, reads can be correctly classified, in principle, if reference genomes from the corresponding taxonomic group are used. In metagenomics experiments, however, it is common that reads from unknown species are sequenced, for which no reference genomes are available from the same species or genus. In this case, reads should be assigned to deeper nodes or edges of the reference tree. This leads to the problem of *phylogenetic placement* of read sequences [4].

Two recent approaches to this task are *pplacer* [26] and *EPA* [4]. These programs are based on probabilistic models of nucleotide substitutions. For a set of *reference genomes*, they require a *multiple alignment* of these genomes – the *reference MSA* –, together with the reference tree. For each query read, *pplacer* evaluates the *posterior probability* of its placement on each edge in the reference tree with regard to the reference *MSA*. It uses several heuristics to speed up the process and has a run time linear with respect to the number of reference taxa, number of queries, and sequence lengths. Similar to *pplacer*, the program *EPA* calculates the likelihood for each query read at each possible edge in the reference tree. *EPA-ng* [3] is a newer version of *EPA* that improves on and greatly accelerates *EPA*.

A more recently developed algorithm for phylogenetic placement of metagenomic reads is *RAPPAS* [23]. As the above mentioned programs, *RAPPAS* needs a reference tree and a reference multiple alignment as input. Unlike these previous methods, however, it does not align the read sequences to the reference sequences or alignment. *RAPPAS* uses so-called *phylo-k-mers*, that are calculated based on the reference tree and alignment in a *preprocessing* step. For each column *i* of the reference alignment, for each edge e of the reference tree and for each possible k-mer *w*, the program calculates the probability to see *w* at the corresponding position *i* in a hypothetical sequence that would branch off from the reference tree at the edge *e*. If this probability is above some threshold, the *k*-mer *w* is called a *phylo-k-mer*. In a preprocessing step, *RAPPAS* assembles a database with all phylo-*k*-mers. Aligning query reads to a reference alignment is a time-consuming step and has, in addition, the potential to introduce errors. The alignment of reads against the reference *MSA* is often performed with *hmmalign* [11, 13]. Alternatively, phylogeny-aware alignment algorithms such as PaPaRa [5] or SEPP [28] can be used.

In contrast to the above three methods, the recently proposed algorithm *APPLES* [2] is a *distance-based* approach. *APPLES* does not require a *MSA* of the reference sequences; in addition, it can take sets of unassembled reads instead of assembled reference genomes as input. The program chooses placement positions based on calculated distances between the reference genomes and the query read sequences. The optimal placement position of a query is the one that minimizes the least squares optimization between the calculated query-reference distances and the distances present in the tree.

In this paper, we present a new approach to phylogenetic placement of metagenomic reads that we call ***A**lignment-free **p**hylogenetic **p**lacement algorithm based on **Spa**ced word **M**atches (App-SpaM). App-SpaM* performs phylogenetic placement without the need for aligned or assembled reference or query sequences. It uses an approach that was originally implemented in the program *Filtered Spaced-Word Matches (FSWM)* [21] to estimate phylogenetic distances between a query and the reference sequences. For two input DNA sequences, *FSWM* estimates the average number of nucleotide substitutions per position, since the two sequences have evolved from their last common ancestor, based on so-called *spaced-word matches*, defined as simple gap-free alignments of a fixed length, with matching nucleotides at certain positions, defined by a binary pattern, and possible mismatches elsewhere. This concept has already been extended to calculate distances between an assembled genome from one species and a set of unassembled reads from a second sequence, or between sets of unassembled reads from two genomes. This adaptation of the program is called *Read-SpaM* [19].

For a set of reference genomes, a reference tree for these genomes and a set of reads from a metagenomics experiment, *App-SpaM* uses spaced-word matches to estimate pairwise distances between every query and every reference sequence. It can then use several fast heuristics to perform phylogenetic placement, based on *phylogenetic distances* calculated from spaced-word matches, and based on the *number* of identified spaced-word matches. In addition to different versions of *App-SpaM* that we implemented, we used the distances calculated from the spaced-word matches as input for *APPLES*.

## Methods

As input, our approach takes a set of *N reference sequences*, an edge-weighted phylogenetic tree *T* – the *reference tree* – with *N* leaves, where each leaf is labelled with one of the reference sequences, and a set of query read sequences. Our algorithm can be divided into three consecutive steps: 1) First, we find so-called *spaced-word matches* between every query and every reference sequence; 2) then we estimate the *phylogenetic distance* between every query and reference sequence based on a ‘filtered’ subset of the identified spaced-word matches 3) at last, we determine the optimal placement position for each query sequence in the reference tree *T*.

### Definitions

For a set Σ of ‘characters’ called the ‘alphabet’, a *sequence over* Σ is an ordered list of elements of Σ. For a sequence *S*, its length is denoted by |*S*|, and for *i* ∈ {1,…, |*S*|}, the *i*-th element of *S* is denoted by *S*[*i*]. The set of sequences of length *n* over Σ is denoted by Σ^*n*^. In the following, we are considering sequences over the set {0, 1} – so-called *patterns* –, over the *nucleotide alphabet* 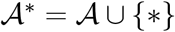 and over the *extended nucleotide alphabet* 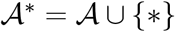. Here, ‘∗’ is a symbol not contained in 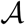, a so-called *wildcard character*.

A spaced word is defined with respect to a given binary pattern *P* ∈ {0,1}^*ℓ*^ of length *ℓ*. A position *j* in the pattern is called a *match* position if *P*[*j*] = 1 and a *don’t care* positions if *P*[*j*] = 0. The number of match positions in a pattern *P* is called the *weight* of *P*. A *spaced word W* over 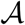 with respect to *P* is defined as a sequence of length |*P*| over 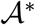, with 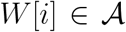 if and only if *i* is a match position of *P*. We say that a spaced word *W* occurs in a sequence *S* over 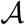 at some position *i* – or that (*W, i*) is a spaced-word occurrence in *S* – if *S*[*i + j* − 1] = *W*[*j*] for all match positions *j* of *P*.

For two sequences *S*_1_ and *S*_2_ and positions *i*_1_ and *i*_2_ in *S*_1_ and S_2_, respectively, we say that there is a *spaced-word match (SpaM)* between *S*_1_ and *S*_2_ at (*i*_1_, *i*_2_), if *S*_1_[*i*_1_ + *j* − 1] = *S*_2_[*i*_2_ + *j* − 1] for all match positions *j* of *P*. In other words, there is a *SpaM* at (*i*_1_, *i*_2_), if there is a spaced word *W*, such that (*W, i*_1_) is a spaced-word occurrence in *S*_1_ and (*W, i*_2_) is a spaced-word occurrence in S2. A spaced-word match with respect to *P* can, thus, be seen as a local gap-free alignment of length |*P*| with matching nucleotides at the match positions of P and possible mismatches at the don’t-care positions, see Fig. 1 for an example. Furthermore, for a substitution matrix assigning a score to any two symbols of the nucleotide alphabet 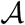, we define the *score* of a spaced word match as the sum of all substitution scores of nucleotide pairs aligned to each other at the don’t care positions of *P*. Spaced-word matches – called *spaced seeds* in this context – have been originally introduced in sequence-database searching [24]; later they were applied in alignment-free sequence comparison, to estimate phylogenetic distances between DNA and protein sequences [30, 21, 20, 32], see [29] for a review.

**Figure 1:**
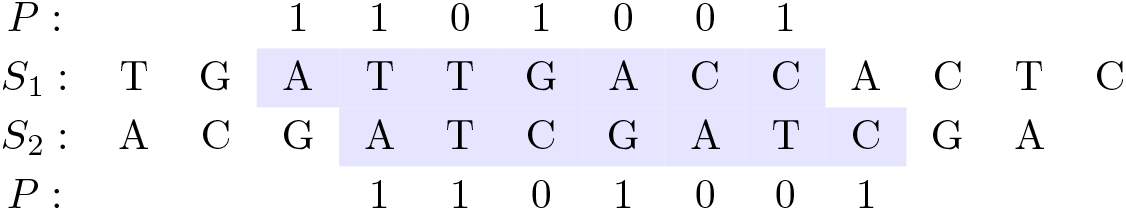
*Spaced-word match (SpaM)* between two DNA sequences *S*_1_ and *S*_2_ at (3, 4) with respect to a binary pattern *P* = 1101001, representing *match positions* (‘1’) and *don’t-care positions* (‘0’). The same spaced word `AT∗G∗∗C` occurs at position 3 in *S*_1_ and at position 4 in *S*_2_. A *SpaM* corresponds to a local gap-free pairwise alignment with matching nucleotides at all *match positions* of *P*, while mismatches are allowed at the *don’t-care* positions.

### Spaced-word matches between query and reference sequences

In a first step, we rapidly identify all spaced-word matches between the query reads and the reference sequences. This can be done by extracting all spaced-word occurrences from the reference and the query sequences and by storing them in two lists *L*_1_ (references) and *L*_2_ (queries) that are both sorted in lexicographic order. Then, for each possible spaced word *W*, all occurrences of *W* in the read and reference sequences would appear as consecutive blocks in the lists *L*_1_ and *L*_2_, respectively.

Once the sorted lists *L*_1_ and *L*_2_ have been established, they can be traversed simultaneously, such that for each spaced word *W*, the blocks with the occurrences of W in *L*_1_ and *L*_2_ are processed at the same time. Each pair of occurrences (*W, i*_1_) in *L*_1_ and (*W, i*_2_) in *L*_2_ corresponds to a spaced-word match at (*i*_1_, *i*_2_) between a reference sequence and a query read. For each such spaced-word match, we calculate its score, and we discard (‘filter out’) all spaced-word matches with a score smaller or equal to some threshold *t*, since low-scoring spaced-word matches can be considered to be *background* or *random* spaced-word matches. As in *FSWM*, the threshold has a default value of *t* = 0, but can be adapted by the user. All spaced words with score larger than *t* are regarded as *homologous* matches.

There is a difference, however, between *App-SpaM* and the original program *FSWM*. In *FSWM*, each spaced-word occurrence (W, i_1_) in sequence S1 can be involved in at most one of the selected (‘filtered’) spaced-word matches. By contrast, a spaced-word occurrence in a read sequence can be matched with multiple spaced-word occurrences in a reference sequence – and vice versa – in *App-SpaM*, as long as the corresponding scores are larger than *t*.

For each query read *Q* and each reference sequence *S*, we store the number *s*(*Q, S*) of spaced-word matches between *Q* and *S* with score larger than *t*. Additionally, we calculate the proportion of mismatches at the don’t care positions of all filtered spaced-word matches between *Q* and *S*, and we estimate the *phylogenetic distance d*(*Q, S*) between *Q* and *S* using the well-known *Jukes-Cantor* formula [18].

In practice, we compute the list *L*_1_ of space-word occurrences from the reference sequences once and hold it in main memory. To limit the number of spaced-word occurrences that are to be held in memory, however, we do not process all spaced-word occurrences from the query reads simultaneously. Instead, we consider a chunk of queries at a time; by default we use chunks 100,000 query sequences. The list *L*_2_ of spaced-word occurrences in the query sequences is then calculated and processed for each chunk separately. This limits the peak usage of main memory while reducing the overhead that we would have if we would process the query reads one-by-one, generating the list *L*_2_ for each query separately.

### Choosing a position for a read in the reference tree

In the following, we propose four heuristics to find a suitable position in our reference tree *T*, where a query read sequence *Q* is added to *T*. For an edge *e* in an edge-weighted tree, let *l*(*e*) denote the length (‘weight’) of *e*. For each query *Q*, we first select an edge *e_Q_* in *T* and insert a new internal node into this edge, thereby splitting *e_Q_* into two new edges *e*_1_ and *e*_2_ with *l*(*e*_1_) + *l*(*e*_2_) = *l*(*e_Q_*). Then, we add a new leaf that is labelled with *Q*, together with a new edge 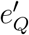 that connects this new leaf with the newly generated internal node. Finally, a length 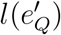 is assigned to the newly generated edge 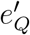.

To find a suitable edge *e_Q_* for a query sequence *Q*, and to assign lengths to the newly generated edges, we are using either the phylogenetic distances *d*(*Q, S*) or the number *s*(*Q, S*) of spaced-word matches with scores larger than *t* between *Q* and all reference sequences *S*. A detailed description how we determine the edge lengths for *e*_1_, *e*_2_, and the newly inserted edge 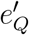 is given in the *supplementary material*. In the rare case that we find no spaced word matches for a query read to any reference sequence, the query is placed at the root. As a fifth approach, in addition to four versions of *App-SpaM*, we used the distance values *d*(*Q, S*) as input for the program *APPLES*.

#### MIN-DIST

In this approach, we first select the reference sequence *S* that minimizes the distance *d*(*Q, S*) over all reference sequences, and we define *e_Q_* to be the edge in *T* that is adjacent to the leaf labelled with *S*. If multiple references have the same smallest distance to *Q*, one of them is chosen randomly.

#### SpaM-COUNT

This works like the previous approach, but instead of selecting the reference sequence *S* that minimizes the distance to *Q*, we select the reference *S* that maximizes the number *s*(*Q, S*) of spaced-word matches with score > *t* between *S* and *Q*.

#### LCA-DIST

Here, we identify the *two* reference sequences *S*_1_ and *S*_2_ with the lowest distances *d*(*Q, S*_1_) and *d*(*Q, S*_2_) to *Q*. Let *v* be the *lowest common ancestor* in *T* of the two leaves that are labelled with *S*_1_ and *S*_2_, respectively. The edge *e_Q_* is then defined as the edge in *T* that connects *v* with its parental node.

#### LCA-COUNT

This is the same as the previous approach, but instead of using reference sequences *S*_1_ and *S*_2_ minimizing the distance with *Q*, we select the two references *S*_1_ and *S*_2_ with the maximal number of spaced-word matches to *Q* with scores larger than *t*.

#### SpaM+APPLES

In addition to the four versions of *App-SpaM*, we used the distances *d*(*Q, S*) between a query *Q* and the reference sequences *S*, calculated as explained above, as input for *APPLES* [2]. *APPLES* performs a least squares optimization to find the position in the tree that fits the calculated distances best. For a reference tree *T*, a query sequence *Q* and input distances between the reference and read sequences, it finds a position for *Q* in *T*, such that the sum of the squared differences between the input distances and the distances in the resulting tree are minimized.

### Evaluation

To evaluate the accuracy and run time of *App-SpaM*, we used the *Placement Evaluation WOrkflows (PEWO)* [22] that have been recently developed to evaluate the accuracy of methods for phylogenetic placement. For a given reference data set, consisting of a set of reference sequences, a reference MSA of these sequences and a reference tree in which the leaves are labelled with the reference sequences, the *pruning-based accuracy evaluation (PAC)* used by *PEWO* evaluates the placement accuracy of an evaluated method as follows: First, a subtree of the reference tree is randomly chosen and removed (‘pruned’) from the reference tree, and the sequences at the leaves of the chosen subtree are removed from the reference MSA. Then, simulated reads are generated by splitting the removed sequences into segments of a given length; these reads are then used as query sequences. An algorithm under evaluation is then used to place the queries onto the reference tree, and the accuracy of the placements is measured.

To measure the accuracy of a placement method, *PEWO* uses the so-called *node distance (ND)*. Here, for each query sequence Q, the distance between proposed placement position of Q and the edge where the subtree was pruned is measured by counting the number of nodes on the corresponding path. The *average* of these distances over all query sequences is then a measure of accuracy for one pruning event. This is repeated with randomly sampled subtrees and *PEWO* uses the *average* accuracy over all sampled subtrees as the overall accuracy of the evaluated method.

*PEWO* also provides a so-called *resources evaluation (RES)* workflow to measure the run time and memory usage of programs. This includes the alignment of queries against the MSA of references for those methods that are based on sequence alignments, or corresponding preprocessing steps, such as the ancestral state reconstruction that is used in *RAPPAS*.

#### Evaluation procedure

We used the *PEWO PAC* workflow to evaluate the placement accuracy of the four versions of *App-SpaM* that we outlined above, namely *MIN-DIST, SpaM-COUNT, LCA-DIST, LCA-COUNT*. In addition, we evaluated the combination of *SpaM* and *APPLES*. In these test runs, we also used different values for the weight *w* in *App-SpaM*, i.e. the number if *match positions* in the underlying pattern *P*.

We also used the *PAC* workflow to compare the accuracy of our approaches to all programs that are included in *PEWO*, using a variety of data sets with different number and length of the reference sequences, and with different degrees of similarity between the reference sequences. At present, the programs *pplacer, EPA, EPA-ng, RAPPAS*, and *APPLES* are included in the *PEWO* package. A quick overview of the reference data sets that we used is given in Tab. 1, a more detailed overview can be found in the supplementary material. For the programs included in *PEWO*, we used multiple parameter combinations as proposed in *PEWO*; a detailed description of these parameters is given in the supplementary materials.

**Table 1:** Data sets used for evaluation. In the following we will refer to each data set by the given *name*. The other columns show the number of reference sequences in the data set, the mean sequence length of these references, and the simulated read lengths used during evaluation. The first eight data sets are used in the *PAC* workflow, the last two in the *RES* workflow. More detailed information is given in the supplementary materials.

| name | number of ref. seq. | mean ref. length | read length |
| --- | --- | --- | --- |
| bac-150 | 150 | 1256 | 150 |
| hiv-104 | 104 | 9096 | 150, 500 |
| neotrop-512 | 512 | 1766 | 150, 300 |
| tara-3748 | 3748 | 1406 | 150, 300 |
| bv-797 | 797 | 1341 | 150 |
| epa-218 | 218 | 1483 | 150 |
| epa-628 | 628 | 780 | 150 |
| epa-714 | 714 | 1169 | 150 |
| CPU-652 | 652 | 1315 | 150 |
| CPU-512 | 512 | 1766 | 150 |

For each reference data set we performed 100 test runs with randomly sampled subtrees and recorded the average *node distance (ND)*. As default, we used a read length of 150 for the simulated queries from the removed references. For some of the data sets we also test additional longer read lengths. To measure the run time and memory efficiency of the evaluated methods, we used *PEWO’s RES* workflow on two data sets of differing sizes, and we recorded the average run time and peak memory usage for different parameter combinations over 5 repeats.

We also performed test runs using (simulated) unassembled sequencing reads as references, instead of contiguous reference sequences. Here, we used different values for the sequencing depth or coverage and compared the placement accuracy with our results on contiguous (‘assembled’) reference sequences. In these experiments, we used the *hiv-104* data set that consists of *HIV* genomes with an average lengths of 9,096 *bp*. We used simulated reads of length 150 and 500, and values for the sequencing depth of 4, 2, 1, 0.5, 0.25, 0.125 and 0.0625.

## Results

Fig. 2 shows the accuracy of the different versions of *App-SpaM*, together with the combination of *SpaM* and *Apples*, with different values of the weight w of the underlying pattern. We used the *bac-150* data set, with a query read length of 150. On average, *MIN-DIST* performed worst and *LCA-COUNT* best among our methods; this is consistent across different values of w. In general, w seems to have little influence on the placement accuracy of *App-SpaM*, but there is no value of w that performs best in all situations. On other data sets we observed comparable results, see supplementary material. The next best placement mode of *App-SpaM* is *LCA-DIST* which is on average 0.77 nodes worse than *LCA-COUNT*, closely followed by *SpaM-COUNT* and *SpaM+APPLES*.

**Figure 2:**
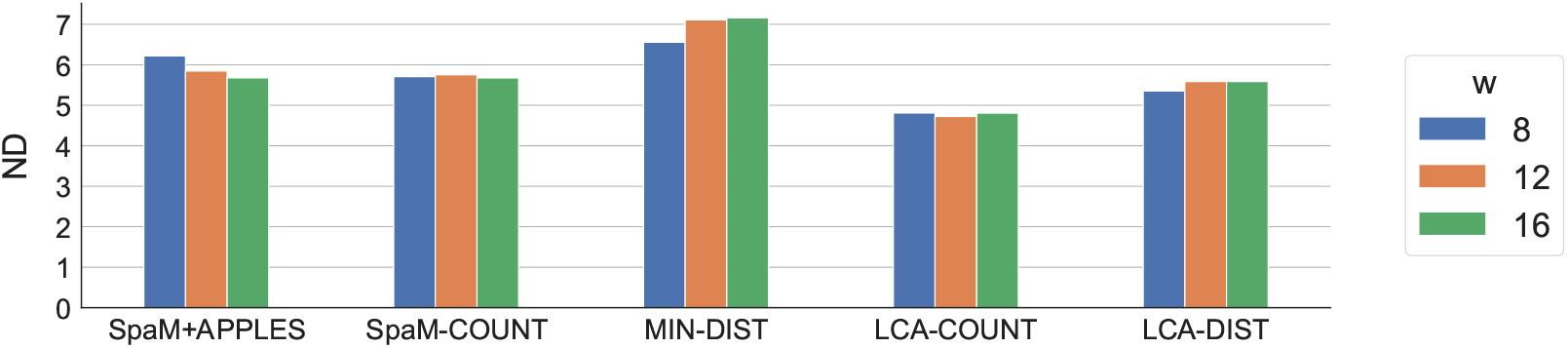
Mean accuracy over *n* = 100 pruning experiments with *PEWO*, measured as *node distance (ND)*, of the four different versions of *App-SpaM* and the combination of *SpaM* and *Apples*. We used three different values, 8, 12 and 16, for the pattern weight *w*, i.e. for the number of *match positions* in the underlying binary pattern.

Fig. 3 compares the performance of *App-SpaM* to five existing placement methods, based on the *node distance (ND)* quality measure, for query read lengths of 150. We used *App-SpaM* with *LCA-COUNT* and with a pattern weight of w = 12. All other programs were ran with the respective optimal parameter settings. Results for other versions of *App-SpaM* and for other parameter settings are given in the supplementary material. As seen in Fig. 3, on most data sets the accuracy of *App-SpaM* is comparable to the accuracy of methods that are based on *Maximum Likelihood*. On average, *pplacer* performed best on the data sets with an average node distance of *ND* = 5.56, followed by *RAPPAS* (*ND* = 5.59), *App-SpaM* (*ND* = 5.7), *EPA* (*ND* = 5.72), *EPA-ng* (*ND* = 5.79), and *APPLES* (*ND* = 9.88). *App-SpaM* performed slightly worse on the data sets with fewer references (*bac-150, hiv-104*, and *epa-218*). On the three data sets that comprise a larger number of reference sequences, our method performed similarly (on *neotrop-512, bv-797*) or slightly better (on *tara-3748, epa-628*) than the competing methods.

**Figure 3:**
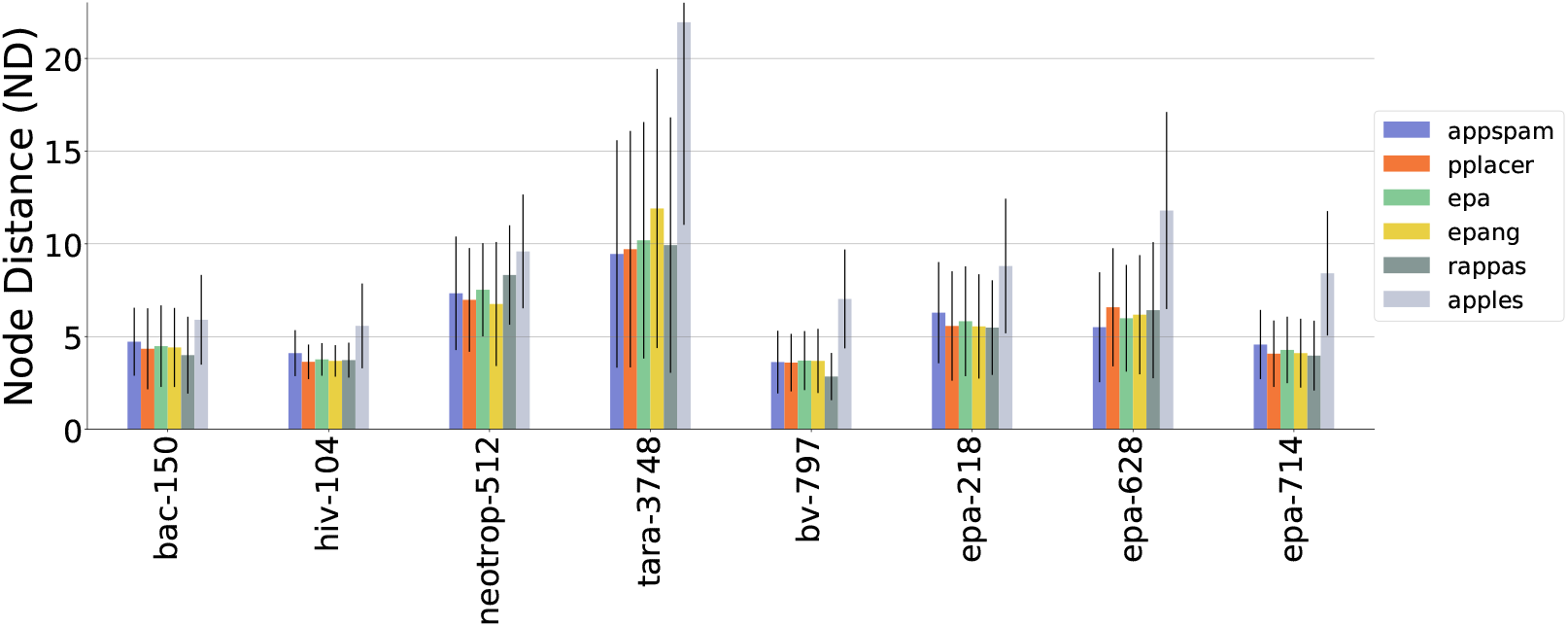
Mean accuracy, measured as in Fig. 2, for *App-SpaM/LCA-COUNT* with a pattern weight *w* = 12, compared to existing placement methods that are included in *PEWO*, on six different reference data sets. Each of the competing methods was run with the respective optimal set of parameter values. Standard deviations across the *n* = 100 test runs are shown as black lines.

Fig. 4 compares the performance of the evaluated methods for two different lengths of the simulated reads that we used as query sequences. Here, we used the data sets *hiv-104, neotrop-512*, and *tara-3748*. As expected, generally all programs are more accurate when longer query reads are used. With a read length of 500, the node distance *ND* improves on average by 0.96 for *hiv-104*, by 1.52 for *neotrop-512*, and by 2.41 for *tara-3748*, compared to a read length of 150.

**Figure 4:**
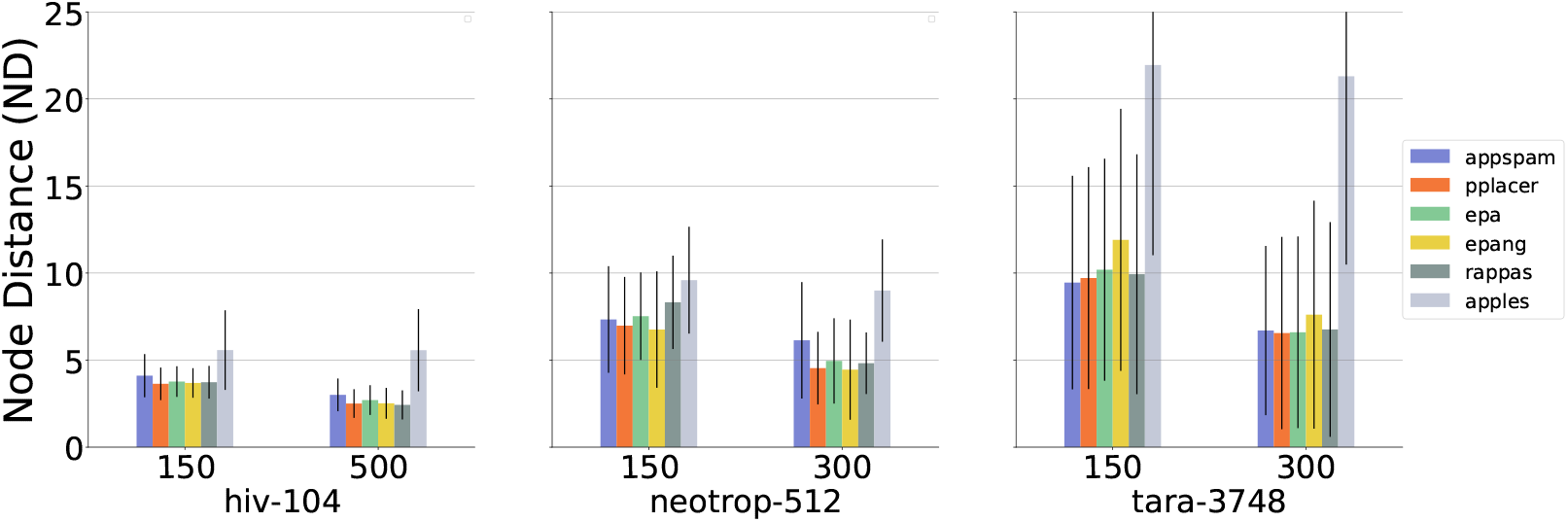
Mean accuracy (*ND*) of best performing parameter combination for each program for different read lengths. Standard deviation across prunings (*n* = 100) shown as black lines.

*App-SpaM* performs best with the placement heuristic *LCA_COUNT*. On average, the best choice for the weight is *w* = 12, but on some data sets, *w* = 8 or *w* = 16 performed slightly better. The number of don’t care positions in the underlying pattern was set to *d* = 32 in all of our test runs. Preliminary results indicate, however, that *d* has little influence on the placement results; we obtained similar results for *d* = 24 and *d* = 32.

*APPLES* performs best with the weighted least squares method *FM* (*k* = 2) and least squares phylogenetic placement (*MLSE*) on nearly all data sets. The performance of *RAPPAS* is strongly dependent on the size of the k used to build the reference data-base of phylo-*k*-mers. Higher values of *k* increase the run times but consistently lead to better results. For all other programs, i.e. *pplacer, EPA*, and *EPA-ng* the performance is nearly identical for all tested parameter settings.

The accuracy of *App-SpaM* on unassembled reference sequences is shown in Fig. 5 for different values of the sequencing coverage, for two different read lengths of the reference sequences (150 and 500), and for different values of the pattern weight w. For a coverage of 1, the *ND* distances are around 1.5 worse than for assembled references. Run time results for two data sets are shown in Tab. 2. We report the run times for preprocessing steps, the placement itself, and the total sum of preprocessing and placement in seconds. Note, that for some programs the run times may vary greatly depending on the chosen parameters. *App-SpaM* with a pattern *weight* of 12 performs 30-60 times faster than the next fastest software with a similar accuracy (*EPA-ng*). Increasing the pattern *weight* further reduces the run time again. When placing multiple queries on the same references, *RAPPAS* average speed will decrease significantly because the time-consuming preprocessing step has to be executed only once.

**Table 2:** Comparison of run times for all tested programs on two datasets. All run times are shown in seconds. For each data set, we show the time for preprocessing (*preproc*.), *placement*, and the *total* sum of preprocessing and placement. Preprocessing includes generating the query alignment or building the phylo-*k*-mer data base for *RAPPAS*. If no parameters are given, the same values as for Fig. 3 were used (see supplementary material).

|  | App-SpaM |  | RAPPAS |  | APPLES | EPA-ng | EPA | pplacer |
| --- | --- | --- | --- | --- | --- | --- | --- | --- |
| | $w = 12$ | $w = 16$ | $k = 6$ | $k = 8$ | | | | |
| CPU-652 |  |  |  |  |  |  |  |  |
| *preproc.* | - | - | 651 | 7253 | 3437 | 3437 | 3437 | 3437 |
| *placement* | 152 | 79 | 710 | 454 | 15226 | 1315 | 194338 | 9257 |
| *total* | 152 | 79 | 1361 | 7707 | 18663 | 4752 | 197775 | 12694 |
| CPU-512 |  |  |  |  |  |  |  |  |
| *preproc.* | - | - | 1070 | 12144 | 1879 | 1879 | 1879 | 1879 |
| *placement* | 34 | 22 | 254 | 185 | 2220 | 127 | 6626 | 1976 |
| *total* | 34 | 22 | 1324 | 12329 | 4099 | 2006 | 8505 | 3855 |

**Figure 5:**
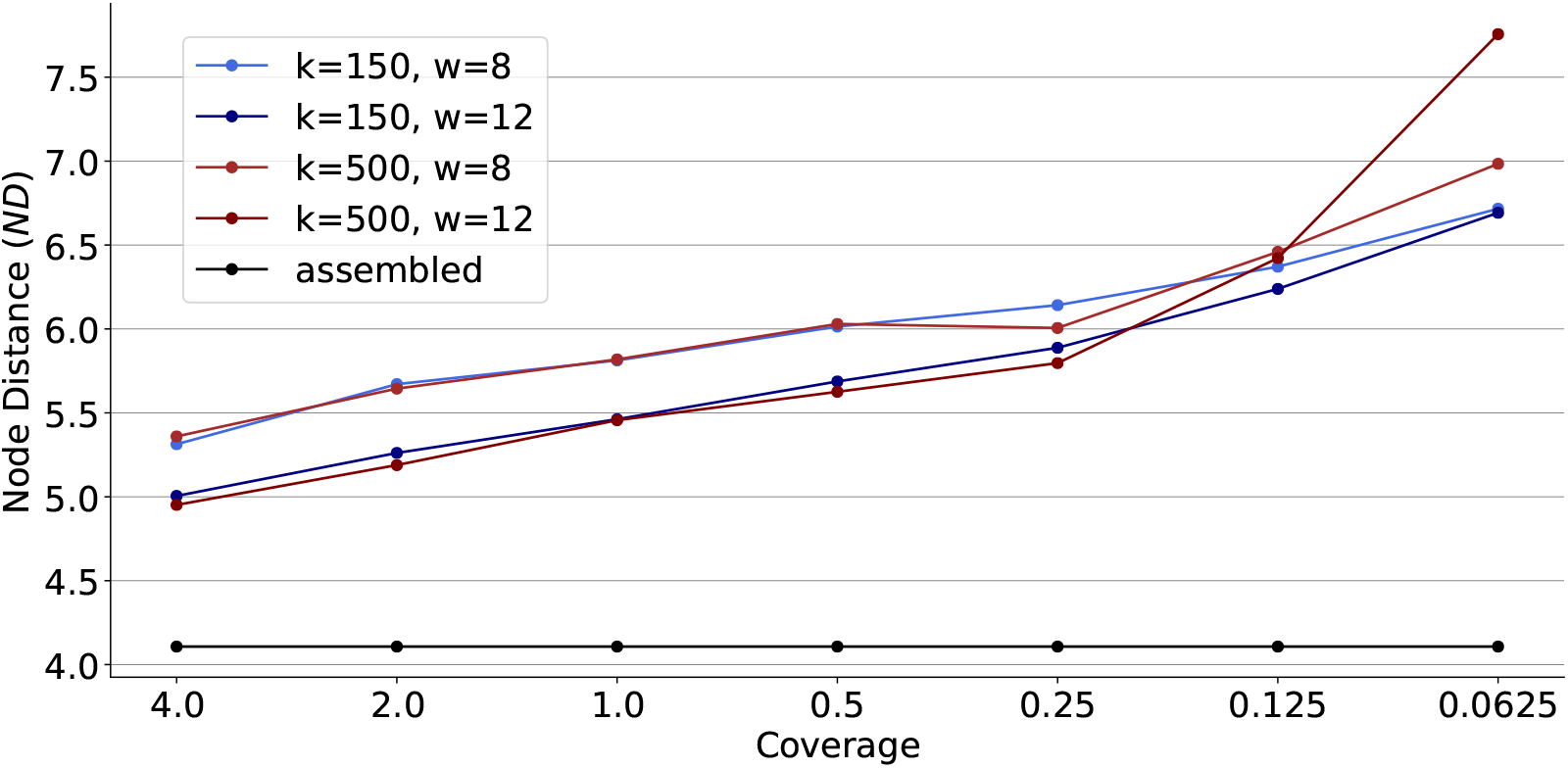
Placement accuracy on unassembled reference sequences of different coverage on the *hiv-104* data set for *App-SpaM* with placement mode *LCA-COUNT*. Results for read lengths of 150 and 500 as well as a pattern *weight* of 8 and 12 are shown in red and blue. The black line serves as reference for the accuracy of *App-SpaM* on the same assembled data.

## Discussion

In this paper, we proposed the new method ***A**lignment-free **p**hylogenetic **p**lacement algorithm based on Spaced word Matches (App-SpaM)* for the task of phylogenetic placement of sequencing reads from metagenomic projects. Phylogenetic placement is the assignment of a query sequence to a position in a reference tree. In contrast to the task of read assignment, where a query read is assigned to one of the reference sequences, phylogenetic placement places reads at arbitrary taxonomic levels and directly incorporates them into the reference tree by inserting new branches and leaves to which the query sequences are assigned. As a result, phylogenetic placement can not only be used to precisely taxonomically classify the reads, but also to update an existing phylogeny one species at a time, as an efficient alternative to de-novo reconstruction of the whole tree.

Existing programs for phylogenetic placement are mostly based on the *maximum likelihood* (ML) principle and therefore require a multiple sequence alignment (MSA) that has to be calculated in a preprocessing step. The query sequences are then aligned against this reference MSA. Both steps are not only time consuming, but also pose limitations on the applicability of phylogenetic placement. At present, only *RAPPAS* does not require a MSA of the reference sequences, and *APPLES* is the only program that is capable of phylogenetic placement without aligned or assembled reference sequences. Our new method *App-SpaM* performs phylogenetic placement without depending on aligned or assembled input sequences. *App-SpaM* reaches nearly the accuracy of alignment-based programs, while it is one to two orders of magnitude faster than existing programs.

*App-SpaM* relies on so-called *spaced-word matches (SpaM)*, simple gap-free local alignments based on a binary pattern of *match* and *don’t-care* positions. Based on these spaced word matches, *App-SpaM* estimates the average number of nucleotide substitutions per position that occurred since two sequences diverged from their last common ancestor in evolution. Spaced-word matches can be utilized to estimate distances not only between assembled genomic sequences, but also between sets of unassembled reads. This gives *App-SpaM* distinct advantages over existing programs: First, the reference sequences do not need to be available as a MSA which is required by all other PP tools except *APPLES*. On its own, calculating the reference MSA is a time-consuming step that can be an additional source of error in the process of phylogenetic placement. Also, some reference sequences might be too long and can not be aligned globally in any meaningful way which completely prevents phylogenetic placement by existing methods. Second, *App-SpaM* does not need assembled reference sequences. It is sufficient to supply a set of unassembled reads for each reference. Fig. 5 shows that *App-SpaM* produces still reasonable results even for low levels of sequencing coverage. Being independent from the availability of assembled references yields additional use cases for phylogenetic placement and greatly simplifies the procedure for all data sets when an approximate estimate of the placement position is sufficient. Third, the preprocessing step of aligning query reads against the reference MSA is omitted in *App-SpaM*, skipping another potentially error-prone procedure. This results in faster run times and a readily accessible software that does not depend on further tools and can be directly executed on the available input data.

Our extensive evaluation showed that *App-SpaM*’s accuracy is close to the accuracy of ML-based methods on most datasets. In some data sets we even achieve comparable placement accuracy with the best programs, but with much shorter run times. The placement heuristic *LCA-COUNT* was clearly the best on all data sets and is therefore used as the default in *App-SpaM*. An important parameter in our spaced-words approach is the *weight* of the underlying binary pattern, i.e. the number of *match positions*. Our program evaluation shows that the results obtained with *App-SpaM* are rather robust for different values of the pattern weight. Several other internal parameters such as the filtering threshold for spaced word matches, the number of don’t care positions, and the number of simultaneously used patterns have default values that perform well for a broad number of data sets and should only be changed with care.

In this study, we implemented a number of simple and fast placement heuristics for determining the query positions within the reference phylogeny. Still, we achieved a high placement accuracy, nearly as good as for the best existing alignment-based ML methods. The placement heuristics that we used, depend either on the number of spaced word matches or the calculated phylogenetic distances between a query read and the references. However, both sources of information might complement one another, potentially leading to overall improved placement results if used properly. Additionally, when using the currently optimal heuristic *LCA-COUNT*, no query placements directly above a reference are possible. This is not only unrealistic but also limits our accuracy performance considerably. Hence, we are continuing to work on additional placement heuristics that use all available information to fully utilize the spaced word matches approach in phylogenetic placement.

The most basic idea would be to calculate a score for every branch in the tree that reflects the likeliness that the query is placed there based on a combined measure of number of spaced word matches and phylogenetic distances to all references below. In theory such a placement method could also be used to express placement uncertainty similar to the likelihood-weight ratios of ML-based methods.

While the run time of *App-SpaM* is already fast, the current implementation is not yet optimized for speed and memory efficiency and multiple strategies to further decrease the run time are possible: First of all, decreasing the run time for large datasets can already be achieved by using a higher weight of the underlying pattern and overall fewer patterns via the available parameter options. Furthermore, all spaced words are completely held in main memory instead of referencing the corresponding positions in the input sequences. This significantly increases the run time and main memory usage and can be improved by more efficient data handling.

Given the test results shown in this study, *App-SpaM* should be a useful tool for one of the most basic tasks in metagenomics data analysis, and efforts should be made to further improve its efficiency.

## Data availability

The source code of *App-SpaM* is freely available through the website *Github*: https://github.com/matthiasblanke/App-SpaM

## Acknowledgments

We thank Benjamin Linard and Nikolai Romashchenko for granting us early access to the PEWO framework and their continued help during the evaluation. We thank Thomas Dencker and Sarah Wendte for helpful discussions on the project. Matthias Blanke was supported by the International Max Planck Research School for Genome Science, part of the Göttingen Graduate Center for Neurosciences, Biophysics, and Molecular Biosciences.

